# Harnessing Extracellular Vesicles for Stabilized and Functional IL-10 Delivery in Macrophage Immunomodulation

**DOI:** 10.1101/2025.01.14.633016

**Authors:** Najla A. Saleh, Matthew A. Gagea, Xheneta Vitija, Tomas Janovic, Jens C. Schmidt, Cheri X. Deng, Masamitsu Kanada

## Abstract

Extracellular vesicles (EVs) are gaining recognition as promising therapeutic carriers for immune modulation. We investigated the potential of EVs derived from HEK293FT cells to stabilize and deliver interleukin-10 (IL-10), a key anti-inflammatory cytokine. Using minicircle (MC) DNA vectors, we achieved IL-10 overexpression and efficient incorporation into EVs, yielding superior stability compared to free, recombinant IL-10 protein. Detailed biophysical and functional analyses revealed that IL-10^+^ EVs contain both monomeric and oligomeric forms of IL-10 on their external surface and encapsulated within vesicles. IL-10^+^ EVs suppressed inflammatory cytokine expression in pro-inflammatory macrophages (from two to 14-fold compared to naïve EVs) without inducing anti-inflammatory polarization, demonstrating a distinct immunosuppressive mechanism. Interestingly, naïve EVs from non-transfected cells also exhibited significant anti-inflammatory effects, suggesting that the intrinsic bioactive cargo of EVs substantially contributes to their function, complicating the interpretation of IL-10-specific effects. Size-based fractionation analyses of IL-10^+^ large EVs (lEVs), small EVs (sEVs), and non-vesicular extracellular particles (NVEPs) revealed IL-10 presence across all fractions, predominantly in monomeric form, with anti-inflammatory activity distributed among subpopulations. Anion exchange chromatography successfully enriched IL-10^+^ exosomes while retaining immunomodulatory effects. However, the shared properties of naïve and IL-10^+^ exosomes underscore the complexity of their immunomodulatory functions. These findings highlight the therapeutic potential of EVs while emphasizing the need to disentangle the contributions of engineered cytokines from endogenous vesicular components.

## INTRODUCTION

Extracellular vesicles (EVs) are emerging as central players in intercellular communication, with significant potential for diagnostics and therapeutics across various diseases, particularly inflammatory and immune-mediated disorders [1]. EVs are categorized based on their biogenesis, size, and composition, with exosomes and ectosomes being the most well-studied subtypes. Exosomes originate from the endosomal pathway, whereas ectosomes bud directly from the plasma membrane, reflecting their diverse formation mechanisms [2–4]. They carry a wide array of cargo, such as proteins, lipids, and nucleic acids, which can modulate recipient cell function upon uptake [2]. As natural carriers of biomolecules, EVs offer unique therapeutic advantages such as high biocompatibility, stability in circulation, and an intrinsic ability to target specific cells or tissues [5]. Recent studies have also demonstrated that EVs can be engineered to display specific protein receptors as molecular decoys, effectively sequestering pro-inflammatory cytokines—a promising approach for treating conditions characterized by immune dysregulation [6, 7]. Besides their potential as engineered delivery platforms, EVs naturally transport various cytokines and other signaling molecules, offering a biological solution that may overcome the fundamental limitations of traditional cytokine-based immunotherapies [8, 9].

Among various cytokines, interleukin-10 (IL-10) has garnered attention for its potent anti-inflammatory effects, making it a promising candidate for immune modulation. IL-10 is crucial in maintaining immune homeostasis, primarily by suppressing pro-inflammatory cytokine production and regulating immune cell function [10]. However, the clinical application of IL-10-based therapies has been significantly limited by the inherent instability and rapid clearance of recombinant IL-10 protein in vivo. These limitations often result in the necessity for high doses to achieve therapeutic outcomes, consequently increasing the risk of adverse effects [11, 12]. Delivering IL-10 protein by EVs offers a potential solution to these challenges, as the vesicular environment may protect IL-10 from degradation and enhance its bioavailability [13, 14]. Studies have shown that vesicular encapsulation can stabilize cytokines, which opens a new therapeutic avenue for improving the pharmacokinetics of immune-modulatory proteins [9].

Macrophages play a central role in regulating immune responses and represent a key target for IL-10-based therapies due to their high expression of IL-10 receptors and tissue localization [15]. These plastic cells can polarize into a pro-inflammatory (M1) or anti-inflammatory (M2) phenotype in response to environmental cues, with M1 macrophages being primary mediators of inflammatory responses [16, 17]. Chronic inflammation is often associated with an imbalance favoring M1 macrophage activity, which can contribute to the pathology of autoimmune and inflammatory diseases [17, 18]. Recent studies have demonstrated that EVs containing cytokines or other anti-inflammatory molecules can selectively modulate macrophage activity, offering a promising therapeutic strategy for conditions such as type 1 diabetes, inflammatory bowel disease, and cardiovascular disorders [19–22].

We used minicircle (MC) DNA vectors to overexpress IL-10 protein and effectively incorporate it into EVs derived from HEK293FT cells. MCs encoding IL-10 allow better transfection efficiencies and prolonged transgene expression with lower cytotoxicity than plasmids [23, 24]. We hypothesized that EVs could effectively stabilize IL-10 and enhance its bioactivity, providing a more robust and durable therapeutic effect than recombinant IL-10 protein alone. Our study evaluated the biophysical and biochemical properties of IL-10^+^ EVs, including their stability, interaction with pro-inflammatory macrophages, and capacity to modulate inflammatory cytokine expression. We observed that overexpressed IL-10 protein was not only incorporated as internal cargo but also highly oligomerized and associated with the external surface of EVs. EV-associated IL-10 protein exhibited improved stability and bioactivity, effectively suppressing pro-inflammatory markers in macrophages. Interestingly, even naïve EVs from non-transfected cells demonstrated significant anti-inflammatory effects, highlighting the complexity of extracellular signaling. These findings underscore how purification methods and the intrinsic heterogeneity of EV populations, including their diverse bioactive cargo and associated non-vesicular extracellular particles, can influence EV-mediated therapeutic delivery.

## METHODS

### Plasmid construct and cloning

All plasmids were constructed using standard PCR cloning methods and verified by sequencing (Genewiz, South Plainfield, NJ). For minicircle (MC) production, *E. coli* strain, ZYCY10P3S2T, and the empty parental plasmid, pMC.BESPX-MCS2, were purchased from System Biosciences (Palo Alto). The cytomegalovirus (CMV) promoter was amplified via PCR and subcloned into pMC.BESPX-MCS2. The DNA template of human IL-10 was synthesized as gBlock (Integrated DNA Technologies) and combined with mScarlet [25] (Addgene plasmid #85042, gift from Dorus Gadella) PCR fragments to generate MC-IL-10-RFP by overlap extension PCR. MC was produced following the established protocol [23]. Briefly, ZYCY10P3S2T was transformed with the parental plasmid and pre-cultured overnight in LB medium containing kanamycin (50 µg/mL). The culture was then expanded overnight in TB medium with kanamycin (50 µg/mL) overnight. MC production was induced by adding equal volumes of LB containing 0.01% L-arabinose and 0.04 mol/L NaOH, followed by 6 h incubation at 32°C. For the EV reporter, pKT2/CAGXSP/PalmReNL was previously developed [26] (Addgene plasmid #182970). Human CD63-GFP (Addgene plasmid #62964, gift from Paul Luzio) was subcloned into the pKT2/CAGXSP vector using recombination cloning (In-Fusion HD Cloning Kit, Clontech), as described before [26, 27]. The DNA templates of human CD9 and CD81 were synthesized as gBlocks fused with GFP by PCR and cloned into pcDNA 3.1 (+) vector (V79020, Invitrogen).

### Cell culture and lentiviral transduction

HEK293FT cells (R700-07, Invitrogen) and RAW 264.7 cells (ATCC) were cultured in Dulbecco’s Modified Eagle Medium (DMEM; Gibco, 11995065) supplemented with 10% (vol/vol) fetal bovine serum (FBS; Cytiva, SH30071.03) and 1% penicillin/streptomycin (Gibco, 15070063). THP-1 cells (kindly provided by Contag Lab) and human peripheral blood mononuclear cells (PBMCs; IQ Biosciences) were cultured in RPMI 1640 Medium (Gibco, 11875093) supplemented with 10% FBS (vol/vol) and 1% penicillin/streptomycin. Primary monocytes were isolated from the PBMCs by magnetic separation using the MojoSort™ Human Pan Monocyte Isolation Kit (BioLegend, 480059) following the manufacturer’s instruction. All cells were incubated at 37°C in a 5% CO_2_ atmosphere. NF-κB reporter THP-1 cells were generated using third-generation lentiviral vectors. These vectors were produced in HEK293FT cells transfected with three plasmids: pHAGE NFkB-TA-LUC-UBC-GFP-W (Addgene plasmid #49343, gift from Darrell Kotton), psPAX2 (Addgene plasmid #12260, gift from Didier Trono), and pMD2.G (Addgene plasmid #12259, gift from Didier Trono) according to the protocol described by Addgene (https://www.addgene.org/protocols/lentivirus-production/) with slight modifications. Plasmid transfection was performed using TransIT-2020 DNA transfection reagent (Mirus Bio, MIR 5400) instead of 25 kDa linear polyethyleneimine (PEI).

### Macrophage differentiation

Macrophages were differentiated from THP-1 cells and PBMC-derived monocytes. THP-1 cells were seeded at a density of 10 × 10⁶ cells/mL and treated with 50 ng/mL phorbol 12-myristate 13-acetate (PMA; Sigma-Aldrich, P8139) for 72 h to induce macrophage-like differentiation. Following PMA treatment, cells were washed twice with phosphate-buffered saline (PBS) to remove residual PMA and incubated in fresh RPMI-1640 medium without PMA for an additional 72 h to allow stabilization. The primary human monocytes were differentiated into macrophages via Macrophage Colony-Stimulating Factor human (M-CSF, 40 ng/mL; Sigma-Aldrich, M6518). To polarize the differentiated cells into an M1-like pro-inflammatory phenotype, cells were further stimulated with 100 ng/mL lipopolysaccharide (LPS; Sigma-Aldrich, LPS25) overnight. Following polarization, M1-like macrophage differentiation was confirmed using qPCR to measure the expression of pro-inflammatory markers, including TNF-α, IL-6, and IL-1β.

### Transfection

HEK293FT cells were transfected with MC-IL10 (with or without an RFP-tag), PalmReNL, CD9-GFP, CD63-GFP, or CD81-GFP using the TransIT-2020 transfection reagent following the manufacturer’s protocol with modification. Briefly, on the same day, 2.5 µg/well of DNA vectors were diluted in 250 µL/well of Opti-MEM (Gibco, 31985062) and gently mixed. Subsequently, 7.5 µL/well of TransIT-2020 reagent was added to the DNA solution, mixed by gentle pipetting, and incubated for 15–20 min at room temperature to allow complex formation. The transfection complex was then added dropwise to each well containing trypsinized HEK293FT cells (2 × 10⁶ cells/well; 6-well plate) in 2.25 mL of complete medium. Following transfection, cells were incubated for 24-48 h before being split into five tissue-culture-treated dishes (100 mm each).

### EV isolation

EVs were isolated from cell supernatant using two different methodologies. For both, EV-depleted FBS was prepared by ultracentrifugation at 100,000 × g, 4°C for 18 h [28]. After transfected and non-transfected (control group) HEK293FT cells reached around 50% confluence, the media were changed to EV-depleted media or serum-free media (without FBS), and the cells were cultured for an additional 72 h to collect the conditioned media. *First isolation method:* small EVs (sEVs)-enriched fractions were isolated as previously described [26] with modifications as described. The conditioned media were centrifuged at 600 × g for 5 min to remove cells and large debris. Subsequently, the supernatants were further filtered through 0.2 µm PES membrane filters (Nalgene, 725-2520) to remove large vesicles. Finally, sEV fractions were collected by a size-based EV isolation method using 50-nm porous membranes (Whatman, WHA110603) with holders (EMD Millipore, SX0002500) by applying a vacuum pressure [29]. The concentrated sEV fractions on the membranes were washed with 5 mL PBS and carefully collected from the membranes. *Second isolation method:* large EV (lEV)-, sEV-, and non-vesicular extracellular particle (NVEP)-enriched fractions were isolated as previously described [30] with modifications. The conditioned media were centrifuged at 2,000 × g for 20 min to remove cells, debris, and apoptotic bodies. The remaining supernatant was centrifuged at 10,000 × g for 30 min to collect the pellet containing lEVs. The pellet of lEVs was washed with PBS at 10,000 × g for 30 min. The supernatant from the first 10,000 × g centrifugation was filtered through 0.2 µm PES membrane filter and ultracentrifuged 130,000 × g for 50 min to collect the pellet containing sEVs. The pellet of sEVs was washed with PBS at 130,000 × g for 50 min. The supernatant from the first 130,000 × g centrifugation was concentrated using Amicon® Ultra Centrifugal Filters, 10 kDa MWCO (Millipore, UFC8010), to reduce the supernatant from 50 mL to 2.5 mL as 10,000 × g for 5 min. The reduced supernatant was washed with PBS to collect the NVEP fraction. To ensure the stability of EVs/NVEPs under freezing conditions, a 10% freezing solution containing 250 mM trehalose, 250 mM HEPES, and 2% bovine serum albumin (BSA) was used as a protective agent [31]. The isolated fractions were aliquoted into 200 µL portions and stored at –80°C until analysis.

### EV purification by anion-exchange column chromatography

The sEVs isolated from the initial methodology underwent additional purification using anion-exchange column chromatography to analyze exosome-enriched fractions. Using a HiTrap DEAE Sepharose Fast Flow column (Cytiva, 17505501), 1 mL of pre-isolated sEVs (from 20 mL conditioned medium) was processed as previously demonstrated with modifications [32]. The anion-exchange column was first equilibrated with an equilibration buffer (10 mM Tris-HCl, pH 7.4, containing 30 mM NaCl). The sEV-containing PBS solution was diluted 5-fold with 10 mM Tris-HCl (pH 7.4) to adjust the NaCl concentration before column loading. After washing with equilibration buffer, EVs bound to DEAE-sepharose were eluted using a linear NaCl gradient (30-500 mM) and collected in 500 µL fractions, 20 fractions in total (10 mL). Electrical conductivity and UV absorbance measurements for each fraction were obtained using an ÄKTA pure HPLC system (GE Life Sciences). Based on UV absorbance profiles, the fractions were pooled into three groups: Protein-high (2 mL), Protein-low (8 mL), and flow-through (10 mL). Each pooled fraction was concentrated to 1 mL using Amicon® Ultra Centrifugal Filters, 100 kDa MWCO (Millipore, UFC5100).

### Nanoparticle tracking analysis

The size distribution and concentration of EVs/NVEPs were determined using Nanoparticle Tracking Analysis (NTA) (ZetaView, Particle Metrix). Samples were diluted in PBS to achieve optimal particle concentration (1 × 10^9^ – 1 × 10^10^ particles/mL) before analysis. Automated measurements were taken at 11 distinct positions in the sample cell, with outlier control to select optimal video quality. Particle size distribution (diameter in nm) and concentration (particles/mL) were calculated based on Brownian motion and particle scattering.

### Protein quantification

Protein concentration was determined using the Micro Bicinchoninic Acid (BCA) Assay (Thermo Fisher Scientific, 23235), following the manufacturer’s protocol. Briefly, the EVs/NVEPs (1 μL) were diluted in 99 μL PBS and incubated with the BCA reagent working solution at 37°C for 2 h. Absorbance was measured at 562 nm using a microplate reader (Spark, Tecan). A standard curve was prepared using bovine serum albumin (BSA) standards ranging from 0.5 to 40 µg/mL. Protein concentrations were calculated by linear regression based on the standard curve.

### Western blotting

The isolated EVs/NVEPs were lysed with 4X sample buffer (Bio-Rad, 1610747) with (for detecting TSG101 and IL-10) or without β-mercaptoethanol (for detecting CD81, CD63, and CD9), ran on a 4-20% Mini-PROTEAN TGX Stain Free gel (Bio-Rad, 4568096) and transferred to PVDF membranes (Millipore, IPFL00010,). The parental cells were also lysed to verify IL-10 and GAPDH. Membranes were blocked with PBS containing 5% skim milk and 0.05% Tween20 (v/v) for 30 min at room temperature; incubated with primary antibodies overnight at 4°C at dilutions recommended by the suppliers as follows: anti-IL-10 (1:5,000; Proteintech, 60269-1-lg), anti-GAPDH (1:10,000; Santa Cruz, sc-365062), anti-TSG101 (1:1,000; Proteintech, 14497), anti-CD81 (1:3,000; Proteintech, 66866), anti-CD9 (1:1,000; Proteintech, 60232), anti-CD63 (1:1,000; Ts63, Thermo Fisher, 10628D). Membranes were washed 3 times with PBS containing 0.05% Tween20 (v/v), incubated with HRP-conjugated anti-mouse (1:10,000; Cell Signaling, 7076) or anti-rabbit (1:10,000; Cell Signaling, 7074) antibodies for 1 h at room temperature, and washed again to remove unbound antibodies. Membranes were visualized with ECL Select Western Blotting Detection Reagent (GE Healthcare, RPN2235) on ChemiDoc MP Imaging System (Bio-Rad).

### Fluorescence microscopy

Fluorescence deconvolved images were taken using the DeltaVision Microscope system (GE Healthcare Life Sciences). Cells (5 × 10^4^) were observed using a glass bottom μ-dish 35 mm (ibidi, 81156). To analyze sEV uptake, 4 × 10^9^ IL-10-RFP^+^ sEVs were incubated with the cells for 17 h. The cells were stained with 10 μg/mL Hoechst 33342 (LifeTechnologies, H3570) before microscopy was performed. The fluorescence filter set for TRITC and DAPI and the 60x oil immersion objective, 1.42 NA, were used to acquire images and process z-stacks (Optical section space: 0.2 µm, Number of optical sections: 30) for deconvolution. Maximum intensity projection images of the z-stack were created using ImageJ software (The National Institutes of Health, Bethesda, MD, USA). To quantify the CD63/CD9/CD81-GFP-positive and IL-10-RFP-positive sEVs, a drop of the isolated sEVs was placed on hydrophobic PTFE printed slides (Electron Microscopy Sciences, 63429-04), as previously described [33]. After 30 min of incubation at 4°C, slides were washed twice with PBS, and images were acquired. Image analysis was conducted using ImageJ/Fiji software [34] with the EVAnalyzer plugin [35]. The ‘EVColoc’ function was used to quantify the number of RFP and GFP-positive sEVs. A statistical analysis was performed on the data produced by EVAnalyzer using an IPython script with thresholds set using the Median ± 1.5*IQR metric.

### Transmission electron microscopy

To analyze the sample morphology, the isolated EVs/NVEPs were fixed in 1% paraformaldehyde. A formvar-coated gold grid was kept in a saturated water environment for 24 h and placed on a 50 μL aliquot of EV solution, allowing it to incubate for 20 min while covered. Next, samples were washed and blocked by placing each one face down on top of a 100 μL droplet of the following solutions: PBS (2×, 3 min), PBS/50 mM Glycine (4×, 3 min), PBS/5% BSA (1×, 10 min). Anti-IL-10 antibody (1:200; Proteintech, 60269-1-lg) in 5% BSA/PBS was used for labeling (1 h), followed by six washes in PBS/0.5% BSA. Samples were incubated in a 1:50 dilution of anti-mouse IgG-gold (Sigma-Aldrich, G6652) in 5% BSA/PBS (20 min) and washed in PBS (6×) and water (6×). The samples were negative stained with 1% uranyl acetate. Excess uranyl acetate was removed by contacting the grid edge with filter paper and subsequently air-dried. Samples were observed using a JEOL 1400 Flash Transmission Electron Microscope equipped with an integrated Matataki Flash sCMOS bottom-mounted camera. The 1400 Flash was operated at 100 kV.

### Proteinase K protection assay

To determine the localization of IL-10 within EVs/NVEPs, aliquots containing 2 × 10^8^ particles were incubated with proteinase K (final concentration 10 µg/mL; Qiagen, 19134) in PBS to a final volume of 50 µL, as previously described [36]. The samples were kept on ice for 20 minutes, followed by treatment with or without 1% Triton X-100 on ice for 10 minutes. The reactions were halted by adding Halt Protease Inhibitor Cocktail (to a final concentration of 1×; Thermo Fisher, 87786). Samples were subsequently prepared for SDS-PAGE and analyzed by Western blotting.

### Binding of human IL-10 recombinant protein (hIL-10 RP) to sEVs

Following established protocols [37], sEVs (2 × 10^9^ particles) or PBS (control group) were incubated with 1 ug/mL of hIL-10 RP (Cell Signaling, 35979) for 2 h at 37°C under constant agitation. Unbound hIL-10 RP was removed by concentrating the sEVs with Amicon® Ultra Centrifugal Filters, 100 kDa MWCO (Millipore Z648043). The concentrated samples were then prepared for SDS-PAGE and analyzed by Western blotting.

### Treatment of sEVs with heparinase II

To investigate the IL-10 protein binding to heparan sulfate, sEVs (2 × 10^9^ particles) were incubated with 40 U/mL of Heparinase II (Sigma-Aldrich, H6512) in a final volume of 200 µL for 2 h at 37°C under agitation, as described in previous studies [38]. Control sEVs were incubated with PBS alone. The reactions were terminated by adding Halt Protease Inhibitor Cocktail to a final concentration of 1×. Samples were then processed for SDS-PAGE and analyzed by Western blotting.

### Inhibition of Glycosaminoglycan (GAG) Synthesis

To inhibit glycosaminoglycan (GAG) synthesis, HEK293 cells transfected with IL-10 were treated with 2.5 mM of the pharmacological inhibitor p-Nitrophenyl-β-D-xylopyranoside (pNP-Xyl; Sigma-Aldrich, 487870) for 72 h, as previously reported [39]. Following treatment, cell supernatants were collected for sEVs isolation and analyzed by Western blotting.

### Bioluminescence assays

The uptake of IL-10^+^ PalmReNL-sEVs and the NF-κB activity in pro-inflammatory M1-like macrophages were analyzed by bioluminescence measurement after incubating the cells with sEVs/NVEPs. Wild-type THP-1 cells or NF-κB-reporter THP-1 cells were plated in 96-well black clear-bottom cell culture plates at a concentration of 50,000 cells/well, with or without polarization to M1-like macrophages via 100 ng/mL LPS stimulation. The culture medium was replaced with an EV-depleted medium or serum-free medium, and IL-10^+^ PalmReNL-sEVs (8 × 10^8^ particles) or IL-10^+^ sEVs/NVEPs (20 μL) were added at the indicated concentrations, with or without pre-incubation with anti-IL-10 receptor alpha antibodies (1 ug/mL; R&D Systems, MAB274). After the incubation period, the cells were washed twice with PBS, and the uptake of reporter sEVs and NF-κB activity were analyzed by measuring bioluminescence signals. Furimazine (25 µM; Promega, N1110) was added for PalmReNL detection, and D-luciferin (30 mg/mL; Promega, E160A) was added for the NF-κB reporter. Assays were performed using a Spark Multimode Microplate Reader (Tecan).

### Quantitative real-time PCR (qPCR)

Total RNA was extracted from isolated cells and EVs/NVEPs using the RNeasy Mini Kit (Qiagen, 74106) and Exosomal RNA Isolation Kit (Norgen Biotek, 58040), respectively, according to the manufacturer’s protocols. RNA concentration and purity were assessed using a NanoDrop spectrophotometer (Thermo Fisher Scientific). cDNA synthesis was performed with 25 ng of RNA using the SuperScrip III Reverse Transcriptase (Invitrogen, 56575) in a 20 µL reaction volume following the manufacturer’s instructions. qPCR was conducted using the SYBR Select Master Mix for CFX (Applied Biosystems, 447942) on a CFX96 Real-Time System (Bio-Rad). Each reaction was performed in triplicate in a total volume of 20 µL, containing 1 µL of diluted cDNA template. Primers were designed to span exon-exon junctions to prevent amplification of genomic DNA (**Supplementary Table 1**). Relative expression levels were calculated using the ΔΔCt method, with normalization to the housekeeping gene GAPDH or CD68. Data analysis was conducted using Bio-Rad CFX Maestro software, and fold changes were calculated relative to the control group. All experiments were conducted in biological triplicates, and data were presented as mean ± standard error.

### Statistical Analysis

All statistical analyses were performed using GraphPad Prism version 10 (GraphPad Software, Inc.). Data are presented as mean ± standard error. For all statistical tests, a p-value < 0.05 was considered statistically significant. Comparisons between the two groups were conducted using an unpaired two-tailed Student’s t-test. For experiments involving comparisons among more than two groups, a one-way analysis of variance (ANOVA) was performed, followed by Tukey’s post-hoc test for multiple comparisons, where appropriate. A two-way ANOVA was applied to assess the effect of two independent variables on a dependent variable. Interaction effects were evaluated, and post-hoc comparisons were performed using Šídák’s correction for multiple testing if a significant interaction was detected. All experiments were conducted with at least three biological replicates, and statistical analysis was carried out on data from independent experiments. Graphical representation of the data was generated using GraphPad Prism, with detailed descriptions provided in figure legends.

## RESULTS

### IL-10 protein associated with small extracellular vesicles (sEVs) is stabilized

To achieve overexpression of human IL-10, we constructed a minicircle DNA vector encoding IL-10 (MC-IL10) with or without a red fluorescent protein (RFP) tag (**Supplementary Fig. 1a**). Following transfection into HEK293FT cells, we observed punctate IL-10-RFP signals within the cytoplasm and near the plasma membrane, suggesting that IL-10 was associated with discrete subcellular structures (**Supplementary Fig. 1b**). Successful overexpression of IL-10 was confirmed at both protein and mRNA levels (**Supplementary Fig. 1c**). We then isolated sEVs from the culture conditioned medium of IL-10-transfected HEK293FT cells. A uniform size distribution, with a peak of approximately 113.3 nm in diameter, of the isolated sEVs was confirmed by nanoparticle tracking analysis (NTA) and transmission electron microscopy (TEM) (**Fig. 1a and Supplementary Fig. 2a**). The presence of IL-10 in the sEVs was validated by protein analysis (**Fig. 1b**), mRNA detection (**Fig. 1c**), and immunogold labeling (**Supplementary Fig. 2a**). Interestingly, IL-10 protein associated with sEVs appeared as multiple bands, indicating the presence of distinct oligomeric forms (**Fig. 1b**). The isolated sEVs were further characterized by EV marker tetraspanins (CD9, CD63, and CD81) (**Fig. 1b**). Single-EV fluorescence microscopy analysis of IL-10-RFP-positive sEVs revealed similar colocalization frequencies with GFP-tagged tetraspanins CD9 (2.8%), CD63 (2.6%), and CD81 (2.1%) (**Supplementary Fig. 2b**). To determine the membrane orientation of sEV-associated IL-10 proteins, we performed a proteinase K protection assay. The results revealed that the monomeric and higher-order oligomeric IL-10 were primarily localized on the surface of sEVs, while the smaller oligomeric form was encapsulated within the vesicles (**Fig. 1d**). Notably, stability assays showed that IL-10 on sEVs exhibited significantly greater stability compared to recombinant human IL-10 (IL-10 RP) when subjected to various stress conditions, including exposure to room temperature (2 h), physiological temperature (37 °C, 2 h), and repeated freeze-thaw cycles (**Fig. 1e).**

**Figure 1.**
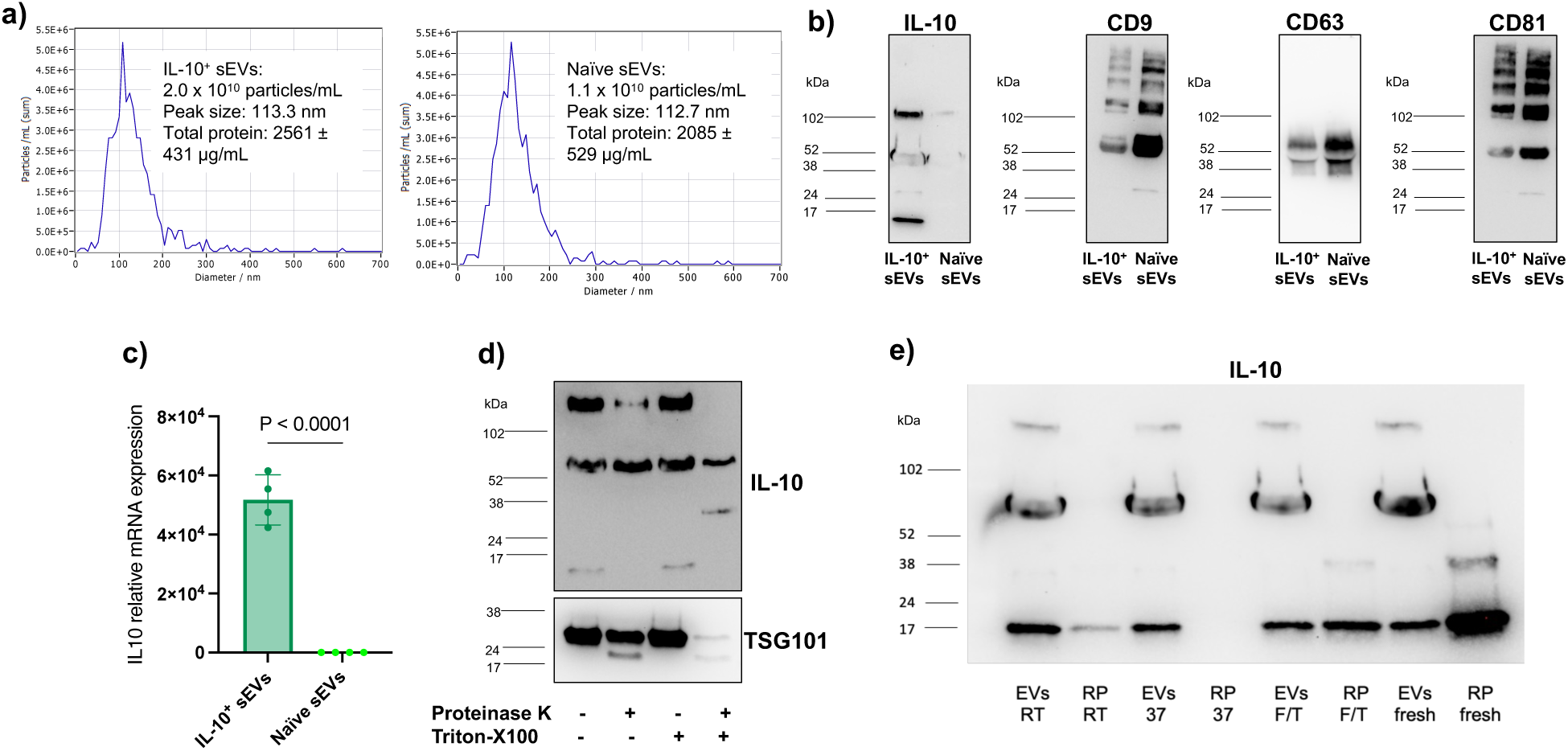
Characterization of IL-10^+^ sEVs. **a)** Size distribution and concentration were measured using nanoparticle tracking analysis. **b)** Western blot analysis of IL-10 protein showing multiple bands and EV markers CD9, CD63, and CD81. **c)** IL-10 mRNA expression analyzed by qPCR. **d)** Proteinase K protection assay for IL-10^+^ sEVs, with TSG101 serving as a luminal protein control. **e)** Stability comparison between sEV-incorporated IL-10 and recombinant human IL-10 protein (RP) under 2 h at room temperature (RT), 2 h at 37 °C (37), and 2 cycles of freezing and thawing (F/T). Fresh samples served as controls. Naïve sEVs were isolated from non-transfected HEK293FT cells.

### IL-10 association with sEVs occurs independently of heparan sulfate proteoglycans (HSPGs) and heparin

To explore the potential interaction between IL-10 and HSPGs on sEVs, we conducted several experimental approaches to investigate whether HSPGs or heparin-mediated interactions influence IL-10 binding to sEVs. HSPGs, known to be present on the surface of cells and EVs, play a critical role in cell signaling, adhesion, and communication [40]. Previous studies have shown that cytokines, including IL-10, may interact with HSPGs, potentially influencing their bioavailability and function in the extracellular space [41]. HEK293FT cells were treated with 4-nitrophenyl β-D-xylopyranoside (PNP-Xyl), an HSPG biosynthesis inhibitor that disrupts the glycosylation of proteoglycans, thereby reducing the availability of functional HSPGs on the cell surface. However, the results were inconclusive due to reduced IL-10 expression in DMSO-treated control cells (**Supplementary Fig. 3a**). This suggests that the treatment conditions may have affected IL-10 production or stability, as low DMSO doses (≥0.25%) have been previously reported to increase pro-inflammatory cytokines [42]. In a second approach, we sought to directly test whether IL-10 binding to sEVs depends on heparan sulfate chains, which are common glycosaminoglycans (GAGs) found on the surface of HSPGs. The heparinase assay demonstrated that IL-10 association with sEVs was independent of heparan sulfate chains, as protein levels remained unchanged post-treatment (**Supplementary Fig. 3b**). To explore further the potential binding of IL-10 to sEVs in a controlled environment, we attempted to synthetically bind recombinant IL-10 to naïve sEVs using a corona formation model [43–46] where proteins or other molecules are expected to adsorb onto the surface of vesicles. However, following a two-hour incubation, we observed no detectable IL-10 monomers on the surface of sEVs (**Supplementary Fig. 3c**). This suggests that IL-10 does not readily bind to the surface of naïve sEVs under these experimental conditions, or if binding occurs, it is weak and transient, making it difficult to detect using the methods employed.

### IL-10-loaded sEVs exhibit anti-inflammatory activity in pro-inflammatory macrophages

To evaluate the cellular uptake of IL-10-loaded sEVs (IL-10^+^ sEVs) by pro-inflammatory macrophages, we employed fluorescence imaging and bioluminescence quantification of reporter sEVs (IL-10-RFP and PalmReNL). Fluorescence microscopy indicated significant uptake of IL-10-RFP-positive sEVs by M1-like macrophages after 17 h, as evidenced by prominent endosomal accumulation of RFP fluorescence signals (**Fig. 2a**). Bioluminescence analysis using PalmReNL-labeled sEVs further demonstrated that IL-10-carrying sEVs achieved three-fold higher uptake compared to control PalmReNL-sEVs (without IL-10) following 2 h of incubation with pro-inflammatory M1-like macrophages (**Fig. 2b**). To investigate the role of IL-10 receptor (IL-10R) in vesicle uptake, we pre-treated macrophages with anti-IL-10Rα blocking antibodies. This treatment resulted in only a modest reduction in bioluminescence signals from IL-10-carrying PalmReNL-sEVs, suggesting that IL-10R interaction is not the primary mechanism for sEV internalization and that receptor-independent uptake pathways are likely involved (**Supplementary Fig. 4a**).

**Figure 2.**
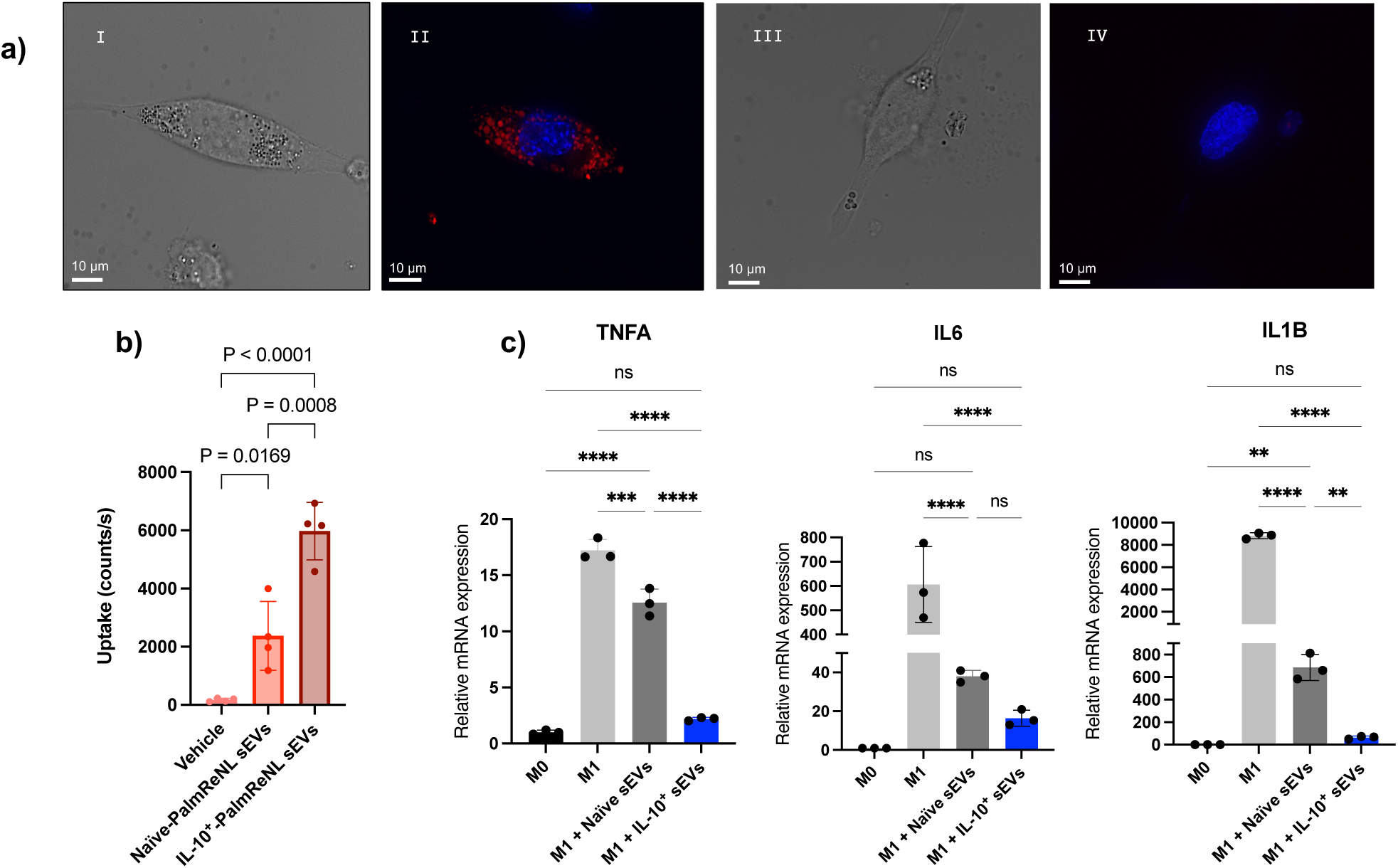
Uptake and functionality of IL-10^+^ sEVs in pro-inflammatory macrophages. **a)** Bright-field (I and III) and fluorescence images (II and IV) showing in red the accumulation of IL-10-RFP^+^ sEVs (I and II) and no sEVs control cells (III and IV) in pro-inflammatory macrophages (THP-1 cells + PMA + LPS) after 17 h incubation. **b)** Uptake of IL-10^+^ PalmReNL-sEVs by pro-inflammatory macrophages assessed by detecting bioluminescence after 2 h incubation. **c)** qPCR analysis of mRNA expression levels of pro-inflammatory markers in LPS-stimulated THP-1 macrophages after 24 h incubation with 10 µg (total protein) of sEVs. M0: non-polarized macrophages; M1: pro-inflammatory macrophages.

Both naïve and IL-10^+^ sEVs demonstrated comparable suppression of NF-κB activity, suggesting shared anti-inflammatory mechanisms (**Supplementary Fig. 4b**). IL-10^+^ sEVs significantly reduced the mRNA expression of pro-inflammatory cytokines TNFA, IL6, and IL1B in pro-inflammatory M1-like macrophages to levels similar to those in non-polarized (M0-like) macrophages (**Fig. 2c**). Notably, IL-10^+^ sEVs did not increase the expression of endogenous anti-inflammatory cytokine genes, including IL-10, indicating that the observed anti-inflammatory effects were primarily mediated by the delivered IL-10 protein (**Supplementary Fig. 5a**).

IL-10^+^ sEVs demonstrated comparable immunomodulatory effects to recombinant IL-10 protein (IL-10-RP) in pro-inflammatory M1-like macrophages. Neither IL-10^+^ sEVs nor IL-10-RP upregulated markers associated with the M2-like anti-inflammatory phenotype of macrophages, such as IL-10, CD163, MRC1, and CD209, suggesting that their effects were primarily immunosuppressive rather than M2-like-polarization (**Supplementary Fig. 5a and Supplementary Fig 5b**). Notably, pre-treatment of IL-10^+^ sEVs with proteinase K revealed that surface-exposed IL-10 selectively reduced TNFA mRNA levels without affecting IL6 or IL1B expression, suggesting IL-10R-mediated modulation of TNFA expression (**Supplementary Fig. 5b**). The anti-inflammatory activity of IL-10^+^ sEVs was validated in primary human macrophages derived from peripheral blood mononuclear cells (PBMCs) (**Supplementary Fig. 6a**). Naïve sEVs exhibited comparable anti-inflammatory effects to IL-10^+^ sEVs in these primary cells. The reduction of pro-inflammatory markers persisted for up to 72 h in both murine (TNFA) and human macrophages (TNFA, IL6, and IL1B) (**Supplementary Fig. 6b and 6c**). However, the potential effects of cell death in macrophages following LPS stimulation cannot be excluded in these prolonged cultures.

### Size-based functional characterization of IL-10-associated EVs

The presence of small particles (smaller than 50 nm) labeled with anti-IL-10 antibodies observed in the TEM images of IL-10^+^ sEVs (**Supplementary Fig. 2a**) prompted us to investigate IL-10-associated EV functionality by size. We fractionated IL-10^+^ EVs into large EVs (lEVs), sEVs, and non-vesicular extracellular particles (NVEPs) using differential ultra-centrifugation and filter concentration (**Fig. 3a**). Protein analysis showed that IL-10 was present in all subpopulations, primarily in monomeric form (**Fig. 3b**). Notably, we often detected weak signals in an oligomeric form in naïve EV fractions, suggesting the existence of endogenous EV-associated IL-10. Functional assays demonstrated that lEVs, sEVs, and NVEPs reduced pro-inflammatory cytokine expression (TNFA, IL6, IL1B) in M1-like pro-inflammatory macrophages, indicating that their anti-inflammatory effects are not size-specific, yet IL-10^+^ NVEPs failed to reduce IL6 expression (**Fig. 3c**). Notably, among the three groups, only IL-10^+^ sEVs showed sustained suppression of NF-κB activity for up to 5 h compared to LPS-only controls, while all groups derived from naïve cells showed reduced anti-inflammatory markers and NF-κB activity, similar to the IL-10^+^ groups (**Fig. 3d**).

**Figure 3.**
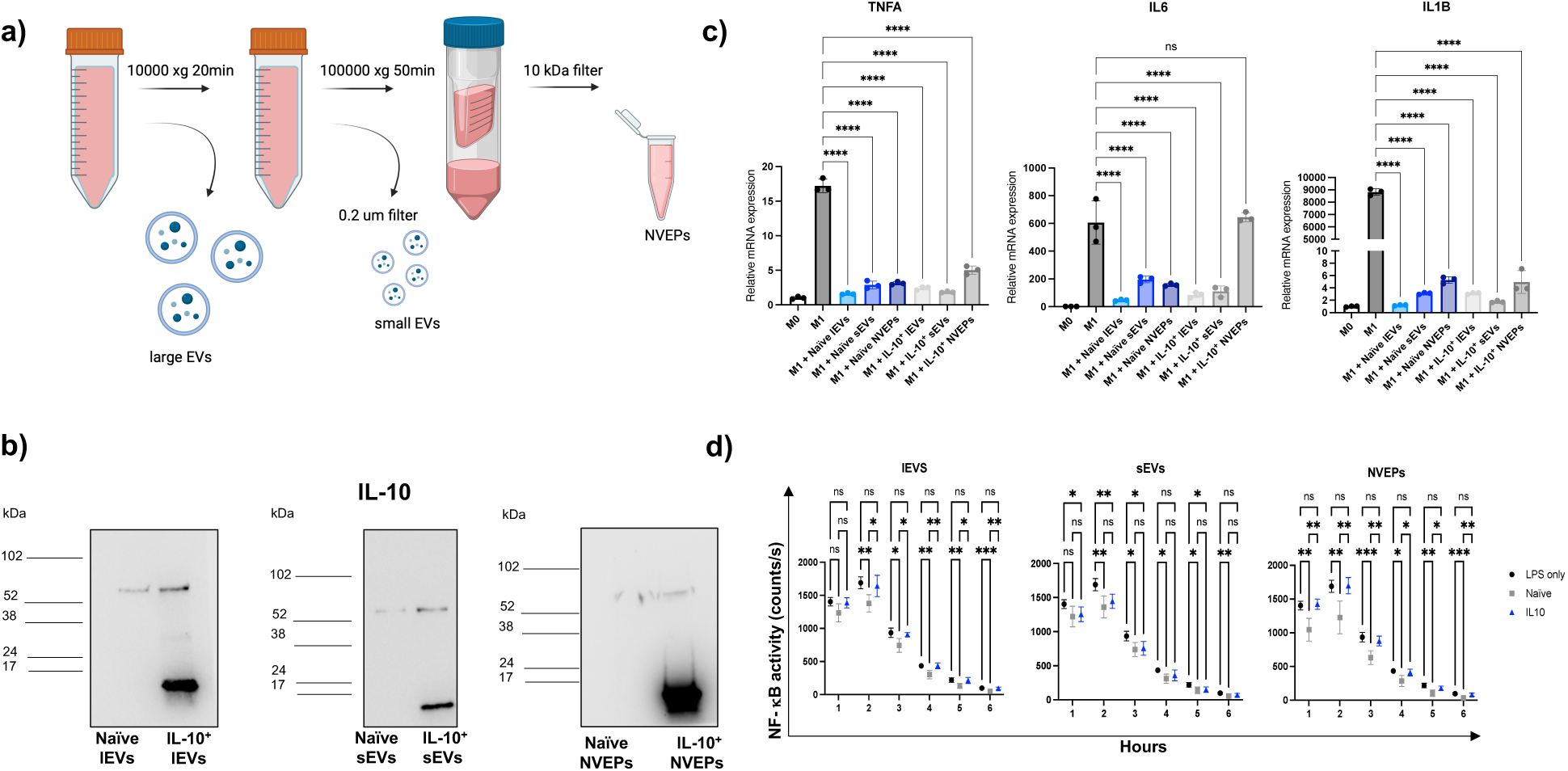
Characterization and functional analysis of IL-10^+^ EVs and NVEPs isolated by differential ultracentrifugation. **a)** Schematic representation of the isolation of large EVs (lEVs) by centrifugation, small EVs (sEVs) by ultracentrifugation (UC), and NVEPs by concentration of UC supernatant (Created with BioRender.com). **b)** Western blot analysis of IL-10 protein expression. **c)** qPCR analysis of mRNA expression of pro-inflammatory markers in LPS-stimulated THP-1 macrophages following 24 h incubation with 50 µL of EVs/NVEPs. **d)** NF-κB activity in M1 macrophages measured by bioluminescence during 1-6 h incubation with EVs/NVEPs.

The presence of EV markers (CD9, CD81) was confirmed in IL-10^+^ EVs/NVEPs, except CD9 in lEVs (**Supplementary Fig. 7a**). TEM revealed the morphology and size of each group (**Supplementary Fig. 7b**). Proteinase K protection assays demonstrated differential IL-10 localization: in lEVs, monomers were present on both internal and external surfaces, while in sEVs, monomers localized primarily on the outer surface. NVEPs contained both IL-10 monomers and oligomers, with only the oligomeric forms showing partial resistance to proteinase K treatment. Complete removal of the monomeric form was observed upon combined treatment with proteinase K and Triton X-100 (**Supplementary Fig. 7c**).

### Protein-high exosomes exhibit an efficient association with IL-10 proteins and anti-inflammatory effects

Anion exchange chromatography was employed to further enhance the purity of IL-10^+^ sEVs as demonstrated previously [32], allowing enrichment of high-performance exosomes based on surface charge using an elution solution with a linear gradient of NaCl (**Fig. 4a**). Three groups of fractions were collected: Protein-high, Protein-low, and Flow-through (**Fig. 4b**). Within the Protein-high group, four fractions exhibited significant enrichment of both IL-10 and exosome markers (CD9, CD63, CD81, and TSG101) (**Fig. 4c**). Western blot analysis revealed that IL-10^+^ exosomes contain three distinct forms of IL-10: monomeric and two oligomeric forms, while naïve exosomes displayed only the lower molecular weight oligomeric form (**Fig. 4d**). Functional assays confirmed that IL-10^+^ exosomes retained their anti-inflammatory effects, effectively reducing pro-inflammatory cytokine expression in M1-like macrophages (**Fig. 4e**). Intriguingly, naïve exosome-enriched fractions showed comparable anti-inflammatory activity in our experimental settings, suggesting that endogenous IL-10 or other bioactive cargo molecules contribute to the observed immunomodulatory signaling mechanisms.

**Figure 4.**
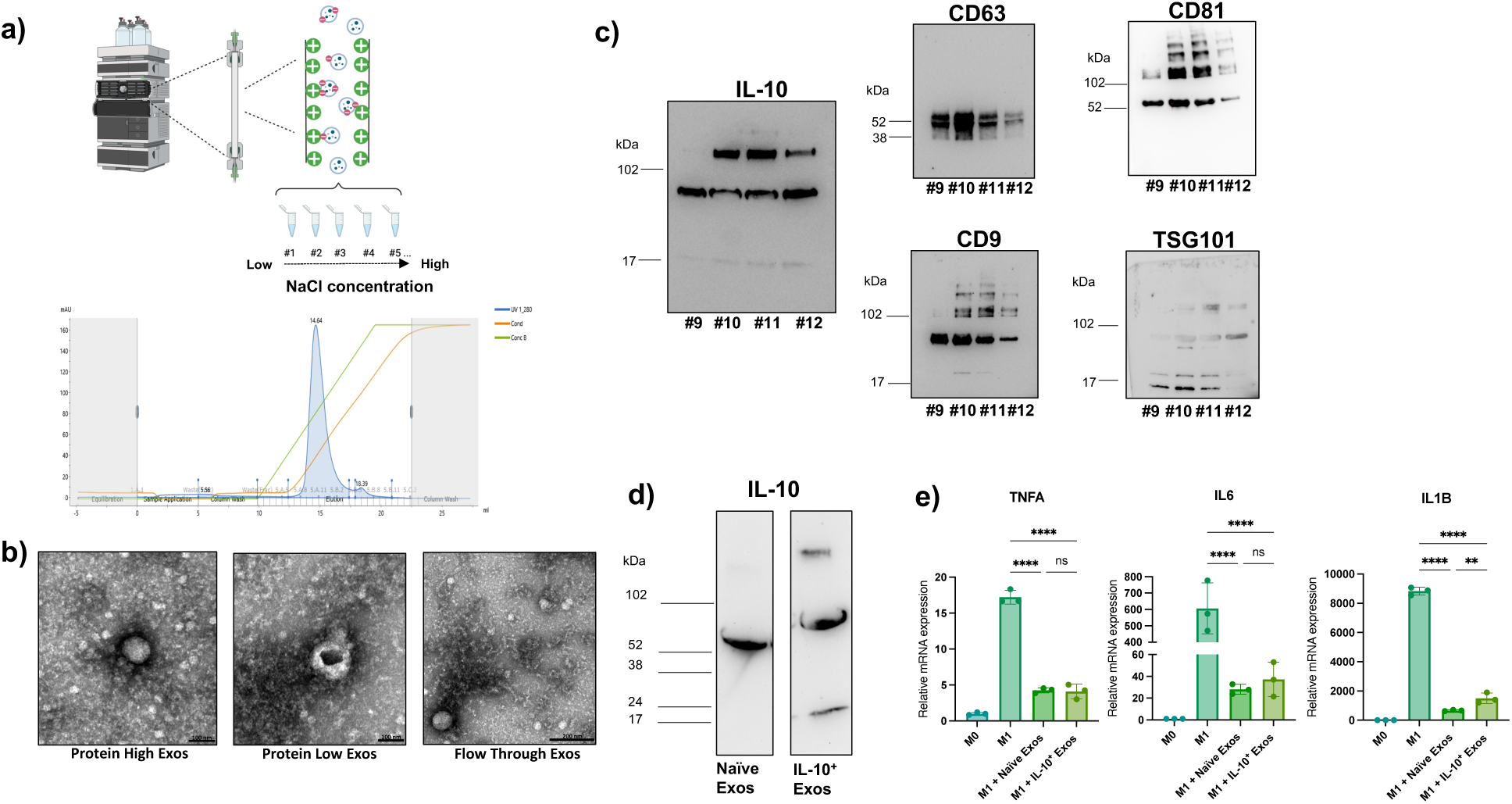
Characterization of IL-10^+^ exosomes (Exos) purified by anion exchange chromatography. **a)** The isolated sEVs underwent a second purification step based on their surface charge by anion exchange chromatography with a linear NaCl gradient (30-500 mM) (Created with BioRender.com). **b)** Representative transmission electron microscopy images of sEVs in Protein-high (left), Protein-low (middle), and Flow-through (right) fractions purified from IL-10^+^ sEVs. **c)** Western blot analysis of IL-10 protein and Exo markers CD9, CD63 and CD81 in Protein-high fractions 9-12. **d)** Western blot analysis comparing IL-10 protein expression between naïve and IL-10^+^ Exos. **e)** qPCR analysis of pro-inflammatory mRNA markers in LPS-stimulated THP-1 macrophages following 24 h of incubation with 10 µg of Exos.

### Intrinsic anti-inflammatory effects of serum-free EVs

Previous studies have highlighted the potential interference of serum-derived EVs and non-vesicular components in investigations of intercellular communication [47, 48]. To exclude potential contributions from serum-derived components to the observed anti-inflammatory activity of naïve sEVs, HEK293FT cells were cultured in serum-free conditions for sEV production. Western blot analysis revealed CD9 was present only in sEVs derived from EV-depleted conditioned media, whereas CD63 was retained in sEVs from both serum-free and EV-depleted conditioned media (**Supplementary Fig. 8a**). This CD63 pattern aligns with previous observation of enhanced CD63 levels under serum-free conditions [48]. Importantly, no significant differences in anti-inflammatory activity were observed between these groups (**Supplementary Fig. 8b**), indicating that the anti-inflammatory properties of sEVs are intrinsic and independent of external serum-derived components.

## DISCUSSION

Building upon recent advances in engineered EVs for therapeutic applications, our study establishes a robust framework for IL-10 stabilization and anti-inflammatory activity using extracellular vesicles (EVs). The results demonstrate that EVs serve as an effective vehicle for IL-10 protein transport, potentially enhancing its therapeutic utility by leveraging the vesicular environment to protect and efficiently deliver IL-10 to target cells.

Our analysis revealed two distinct oligomeric forms of IL-10 proteins associated with sEVs, in addition to the monomeric form. The presence of monomeric and oligomeric IL-10 in our sEV preparations suggests that these vesicles may provide unique structural stabilization of the cytokine, particularly given that multimerization and structural modifications are critical for IL-10 stability [11, 49]. Besides IL-10 oligomers found in sEVs lumen, IL-10 monomers were efficiently retained on the external surface of sEVs, suggesting a corona formation might play a role. Recent studies reveal that molecules such as DNA and proteins can be adsorbed from the extracellular environment onto the outer surface of EVs, creating an EV surface corona [43–46]. This corona acts as a dynamic interface for communication between EVs and their surroundings [50]. For instance, proteins from blood plasma and immunomodulatory and pro-angiogenic proteins may adsorb onto circulating EVs, modifying their signaling functions [43–46]. However, in our experiment, we could not reproduce this effect by artificially decorating EV surfaces with recombinant IL-10 protein (**Supplementary Fig. 3c**). This result suggests that the formation of IL-10-containing protein corona may form naturally and spontaneously during biogenesis [51]. Previous studies have demonstrated that cytokines commonly bind to glycosaminoglycan (GAG) side chains of proteoglycans on EV surfaces [40], with these cytokine-GAG interactions exhibiting remarkable specificity even in the absence of cytokine receptors on EVs [37]. Our heparan sulfate binding assay, however, did not detect IL-10 binding under our experimental conditions (**Supplementary Fig. 3b**). Nonetheless, we cannot rule out the possibility that IL-10 binds to other GAG chains, such as chondroitin sulfate or keratan sulfate. Given the complexity of EV cargo loading and potential receptor-mediated interactions, further investigation is required to identify the specific molecular mechanisms governing IL-10 association with EVs.

IL-10^+^ sEVs exhibited significant uptake and anti-inflammatory effects in M1-like macrophages, as demonstrated by a marked reduction in key pro-inflammatory cytokines (TNFA, IL6, and IL1B), bringing their levels closer to those observed in M0-like macrophages and sustained reduction of NF-κB activity in pro-inflammatory macrophages. Reducing inflammatory markers in primary macrophages reinforces that the effect is not limited to immortalized cell lines such as THP-1 cells. Our future research will test other immune cells, such as neutrophils and regulatory T cells, to advance our understanding of EV-mediated immunoregulation. Unlike other IL-10 studies [14], we did not observe the re-polarization of M1-like macrophages to an M2-like phenotype, suggesting that the observed effects were immunosuppressive rather than polarizing. This discrepancy may be due to differences between in vitro and in vivo conditions, as tissue environments facilitate complex interactions among diverse immune cells, which may promote M1 to M2 macrophage repolarization. Notably, IL-10 on the surface of sEVs selectively downregulated TNFA expression without significantly affecting IL6 or IL1B levels (**Supplementary Fig. 5b**), suggesting a receptor-mediated interaction through the IL-10 receptor that reduces NF-κB activity. This selective effect aligns with previously reported IL-10-mediated anti-inflammatory pathways [52–55]. Additionally, a novel mechanism may involve the delivered IL-10 protein that initiates signaling within endosomes, similar to the recently reported EV-mediated TGFβ signaling in endosomes [39]. Alternatively, luminal IL-10 protein within EVs could be re-released to mediate signaling through autocrine or paracrine pathways [56]. Recent findings using bioluminescent EV reporters revealed that EV cargo could be re-released from recipient cells [57]. Similarly, another study demonstrated that EVs derived from cancer cells encapsulated VEGF and uniquely transported it into the nucleus, where it interacts with nuclear VEGF receptors [58].

Interestingly, size-based fractionation of EVs and NVEPs by sequential ultracentrifugation combined with filter concentration [30, 59] revealed that IL-10’s anti-inflammatory effects are not exclusively dependent on their sizes. All fractions contained IL-10 and exhibited similar reductions in inflammatory cytokine expression. However, IL-10^+^ sEVs uniquely showed sustained NF-κB suppression in pro-inflammatory macrophages, suggesting a potential functional advantage specific to sEVs (**Fig. 3d**). Proteinase K protection assays indicated that IL-10 monomers were primarily located on the external surface of both small and large EVs. In contrast, NVEPs contained both monomeric and oligomeric IL-10 forms (**Supplementary Fig. 7c**), suggesting structural heterogeneity that may influence cytokine release and mechanisms of cellular interaction. Additionally, our findings show that the isolation process involving serial centrifugations contributes to the loss of the higher-order oligomeric form of IL-10 (≥ 102 kDa), highlighting its sensitivity to this methodology. Previous studies have demonstrated that high shear forces during ultracentrifugation can potentially disrupt EV structure [60].

Enhanced purification of IL-10^+^ sEVs using anion exchange chromatography, as recently demonstrated [32], confirmed that exosome enrichment based on surface charge and protein-high properties effectively isolates IL-10-enriched exosomes with potent anti-inflammatory functionality. Notably, exosome-enriched fractions derived from naïve and IL-10-transfected HEK293FT cells contained lower molecular weight oligomeric IL-10, while monomeric and higher-order oligomers were only detected in protein-high fractions isolated from IL-10-transfected cells. Despite these molecular differences, both exosome-enriched fractions exhibited comparable anti-inflammatory activities in pro-inflammatory macrophages.

Finally, we isolated EVs under serum-free conditions, as recently demonstrated [47, 48], to explore the potential contribution of culture conditions and serum contamination to the anti-inflammatory effects of naïve EV fractions. When comparing sEVs isolated from serum-free medium to those obtained from EV-depleted medium, the former expressed CD63 but not CD9. Their modulation of NF-κB activity in THP-1 cells showed no correlation with serum-derived components [61]. The intrinsic immunomodulatory properties of naïve sEVs observed in this study add further complexity to their therapeutic potential. Their anti-inflammatory activity may arise from endogenous bioactive molecules or the inherent structural features of these vesicles. Anti-inflammatory characteristics are also observed in EVs derived from various cell types, such as neural, mesenchymal, and embryonic stem cells [62–64]. Our findings underscore the need for further investigation into endogenous molecular compositions of naïve EVs, which could be leveraged to complement or enhance the efficacy of engineered vesicles. Recent in vivo studies have demonstrated functional IL-10 delivery by EVs to inhibit pro-inflammatory cytokine production in mouse models [14, 56, 65]. Building on these findings, future research should delve deeper into EV-mediated immune signaling mechanisms, including the processes of IL-10 incorporation and release within EVs and NVEPs. Identifying additional bioactive cargo molecules that contribute to their immunomodulatory effects and engineering EVs to deliver multiple therapeutic agents simultaneously are also crucial research areas. Further research focusing on the targeted delivery of EVs to specific macrophage subpopulations or other immune cells, as well as evaluating their therapeutic potential in autoimmune and chronic inflammatory disease models, is highly desired.

## ACKNOWLEDGMENT

We thank Dr. M. Bernard at the MSU Flow Cytometry Core Facility, Dr. A. Withrow at the Center for Advanced Microscopy at MSU, Dr. S. Makaremi at the IQ Microscopy Core Facility, Dr. A. Gilad and lab members, and Dr. G. Perez for their generous support of our daily experiments. This study was supported, in part, by funding from the NIH: R21-EB033554, R01-EB030565, R01-EY016077, and start-up funds from Michigan State University.

## Supplemental Information

**Supplementary Table 1.**
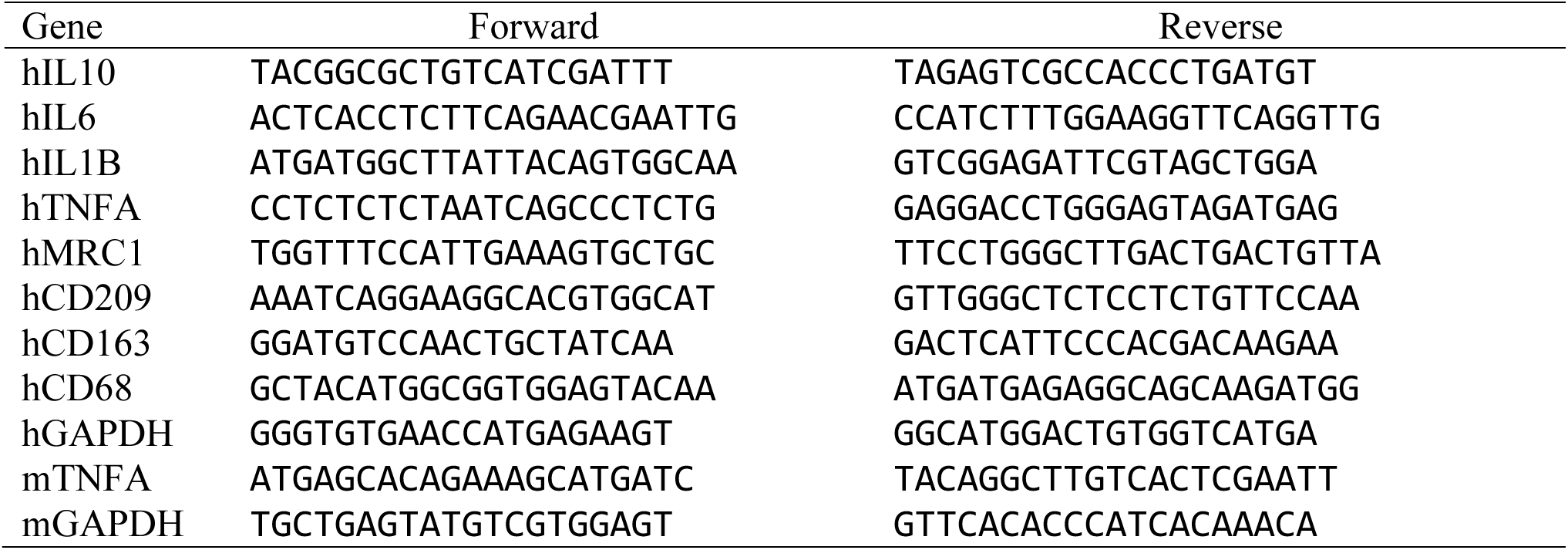
List of primers used in this study. h: Human; m: Mouse.

**Supplementary Figure 1.**
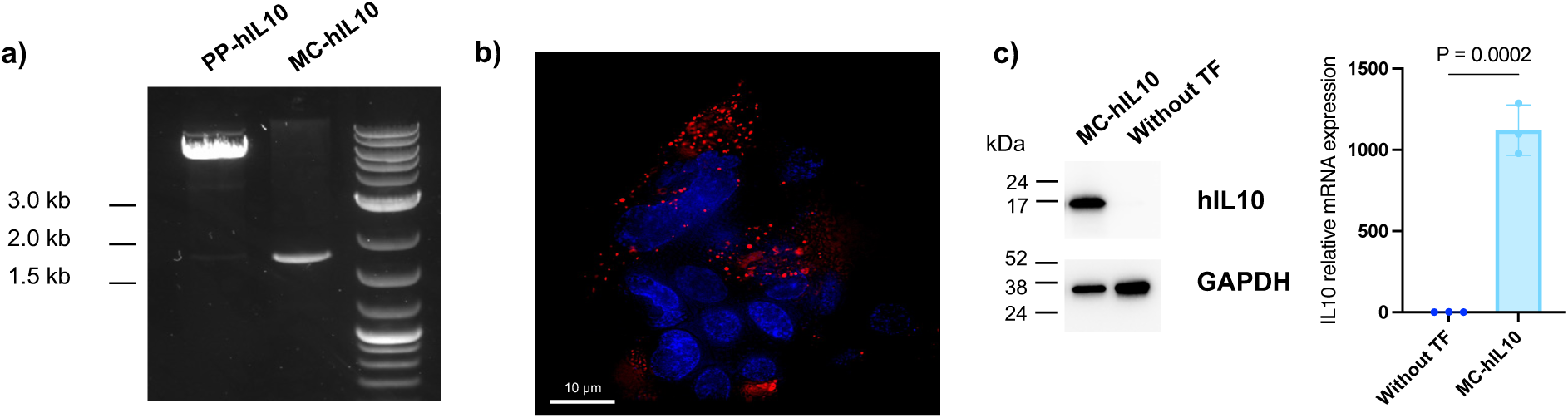
Minicircle-IL10 (MC-IL10) production and transfection. **a)** Agarose gel electrophoresis confirms the ability to generate parental plasmid (PP)– and MC-IL10. DNAs were digested with HindIII. **b)** HEK293FT cells transfected with MC-IL10-RFP. Fluorescence signals of RFP (red; mScarlet) were merged with nuclei stained with Hoechst 33342 (blue). **c)** IL10 expression was confirmed by western blotting and qPCR, while IL10-protein and –mRNA were not detected in HEK293FT cells without transfection (TF). GAPDH indicates a loading control.

**Supplementary Figure 2.**
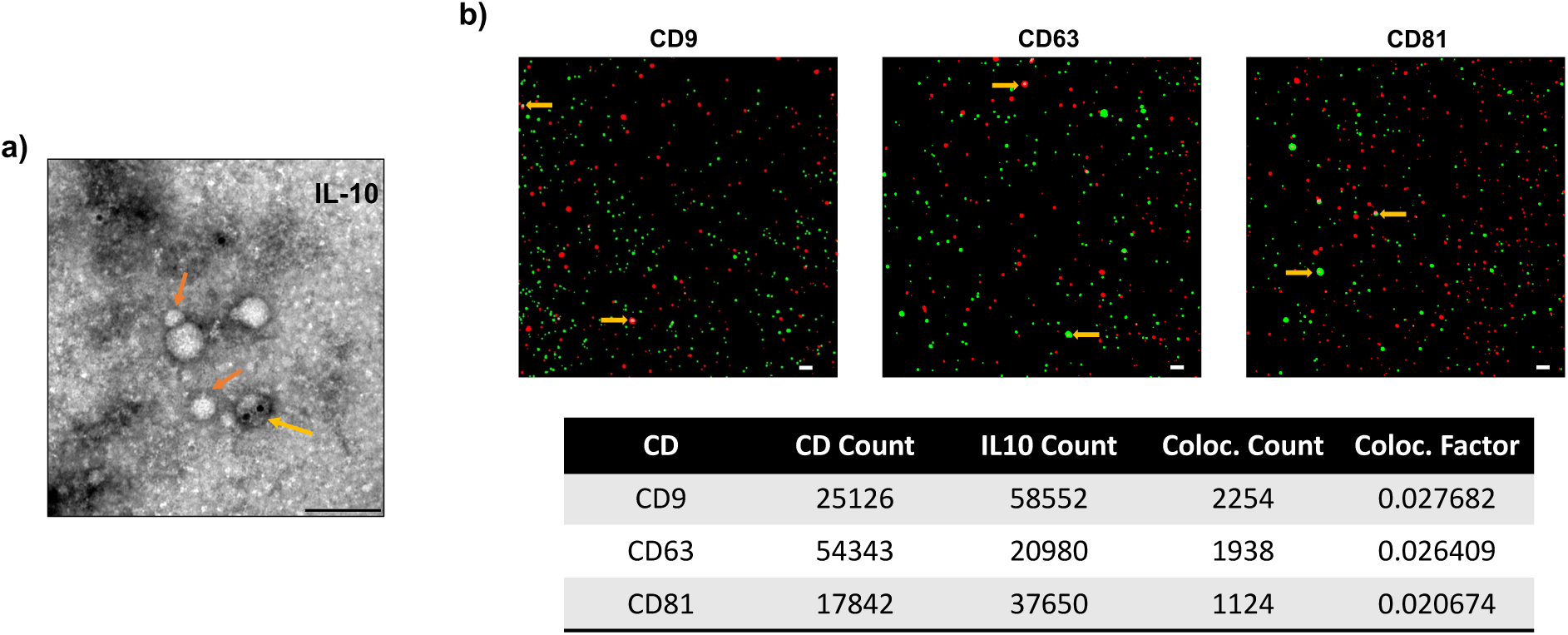
RFP-based labeling of EVs. **a)** Transmission electron microscopy of HEK293FT cell-derived IL-10^+^ PalmReNL-sEVs immunogold labeled for IL-10 (yellow arrow) and small particles (< 50 nm – orange arrows). Scale bar = 100 nm. **b)** Single-EV analysis using fluorescent microscopy to characterize the co-localization (yellow arrows) of IL-10-RFP^+^ sEVs (red) with EV marker tetraspanins—CD9-GFP, CD63-GFP, or CD81-GFP (green). Scale bar = 1 μm.

**Supplementary Figure 3.**
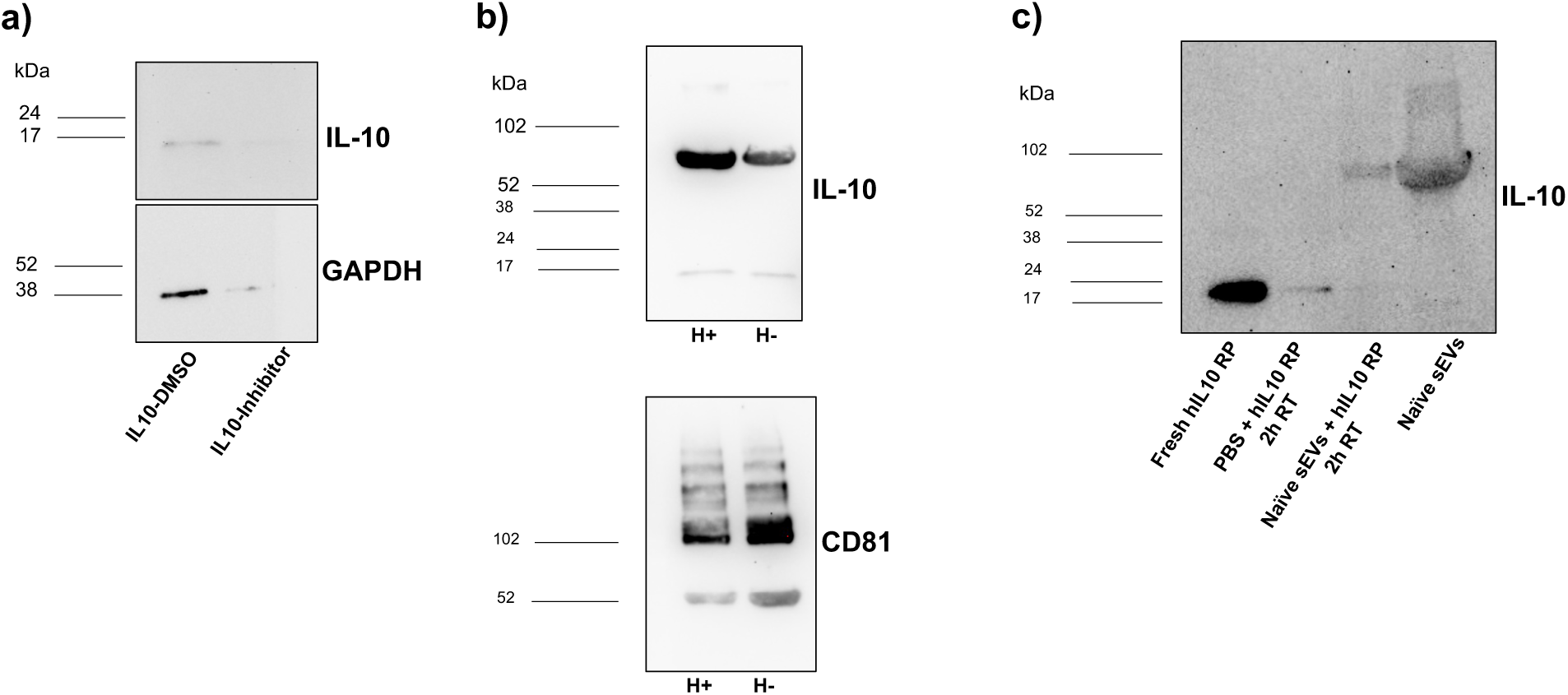
Investigation of IL-10 binding on cells and sEVs. **a-c**) IL-10 protein expression by Western blotting on **(a)** cells after incubation with the inhibitor of heparan sulfate proteoglycan (HSPG) biosynthesis, 4-nitrophenyl β-D-xylopyranoside (PNP-Xyl), **(b)** IL-10^+^ sEVs after incubation with heparinase (H+) and **(c)** naïve sEVs after incubation with human IL-10 recombinant protein (hIL10 RP) at room temperature (RT).

**Supplementary Figure 4.**
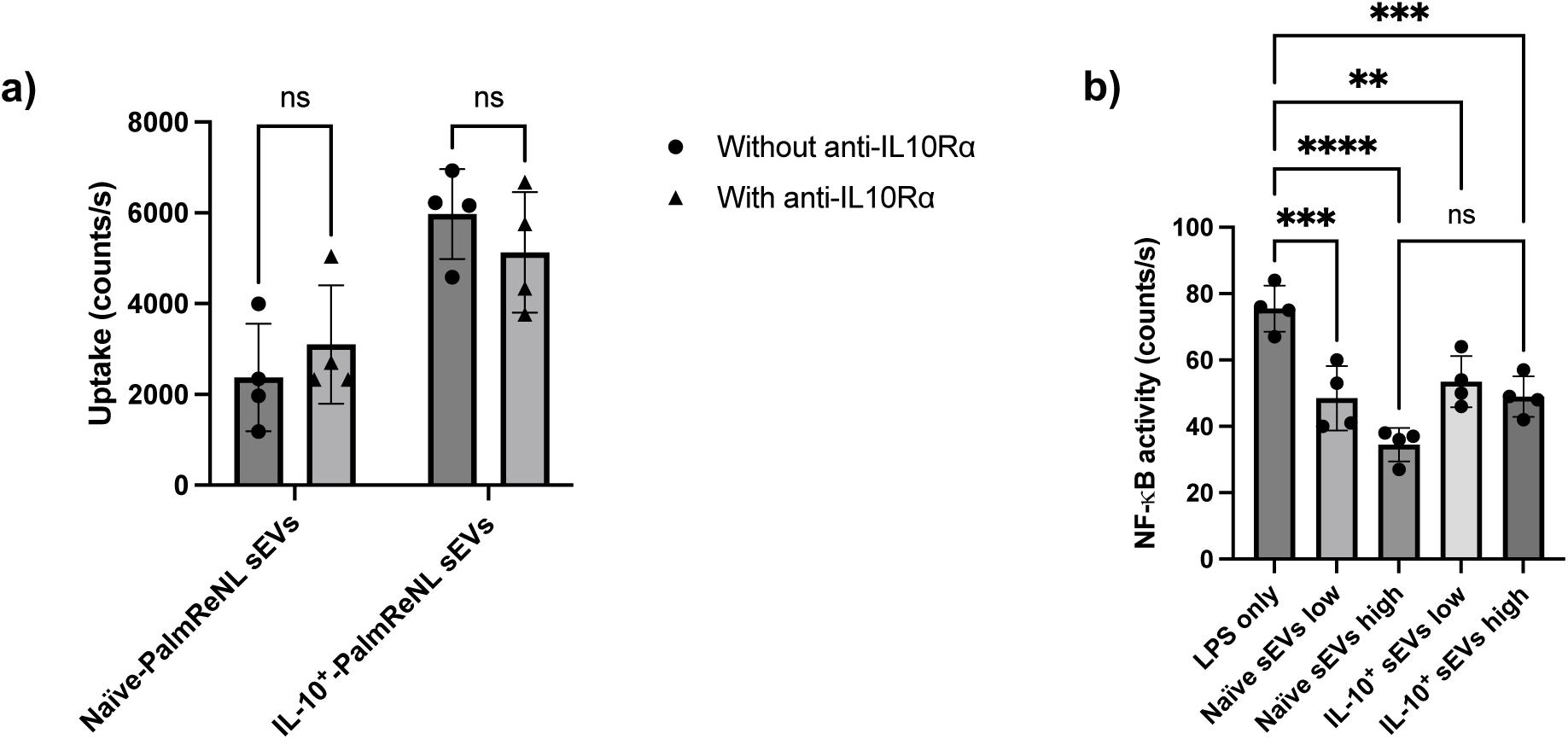
IL-10^+^ sEVs uptake and functionality. **a)** Uptake of PalmReNL sEVs by M1 macrophages after incubation with anti-IL10 receptor alpha antibodies (anti-IL10Rα). **b)** Bioluminescence evaluation of NF-κB activity on monocytes after incubation with sEVs in a low (2 µg) and high (20 µg) concentration.

**Supplementary Figure 5.**
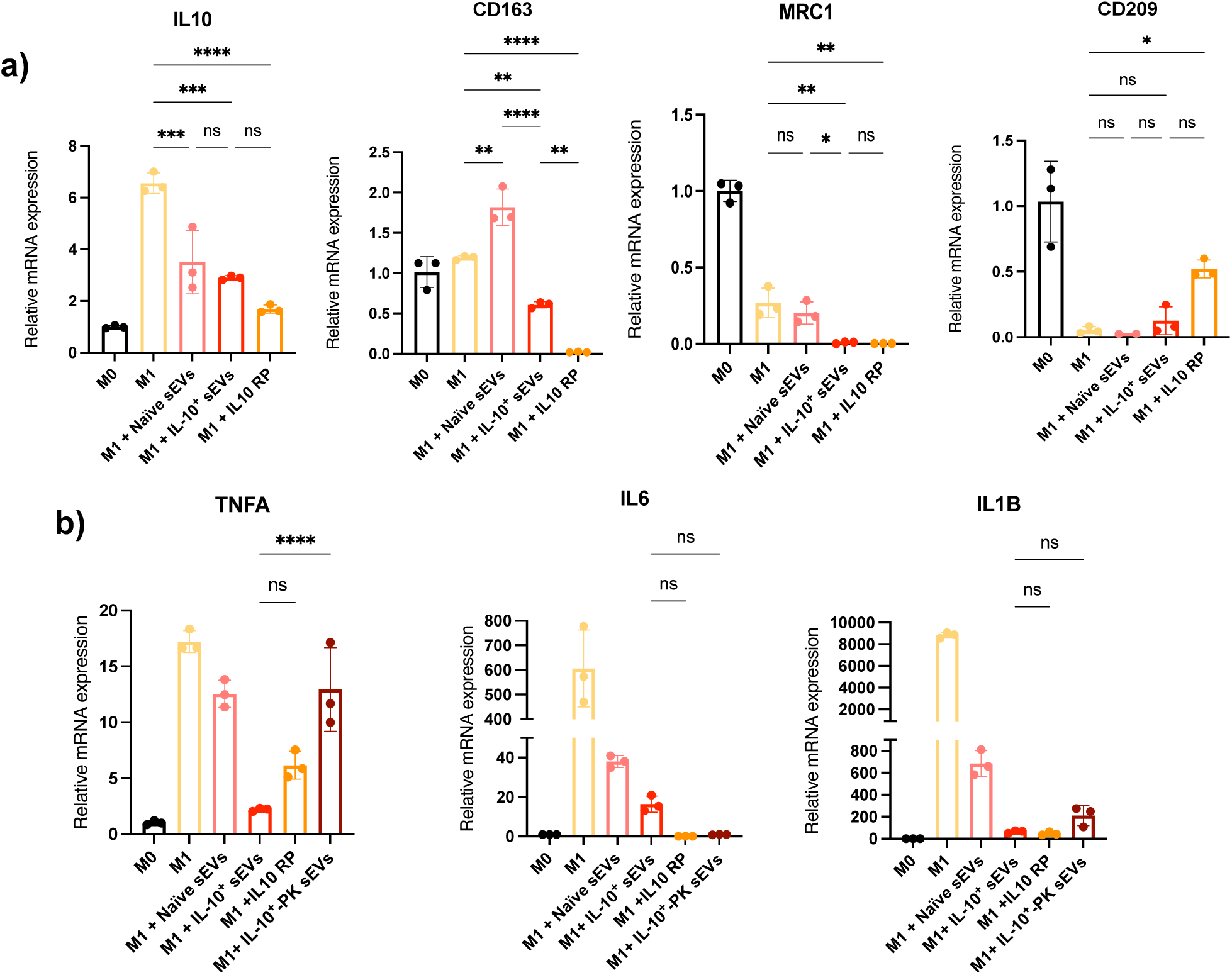
Effect of recombinant IL-10 protein and IL-10^+^ sEVs pre-treated with proteinase K (PK) on inflammatory macrophages. qPCR analysis of **(a)** anti-inflammatory and **(b)** pro-inflammatory marker mRNA expression in LPS-stimulated THP-1 macrophages following 24 h incubation with sEVs (10 µg total protein) and recombinant IL-10 protein (IL-10 RP; 10 ng/mL).

**Supplementary Figure 6.**
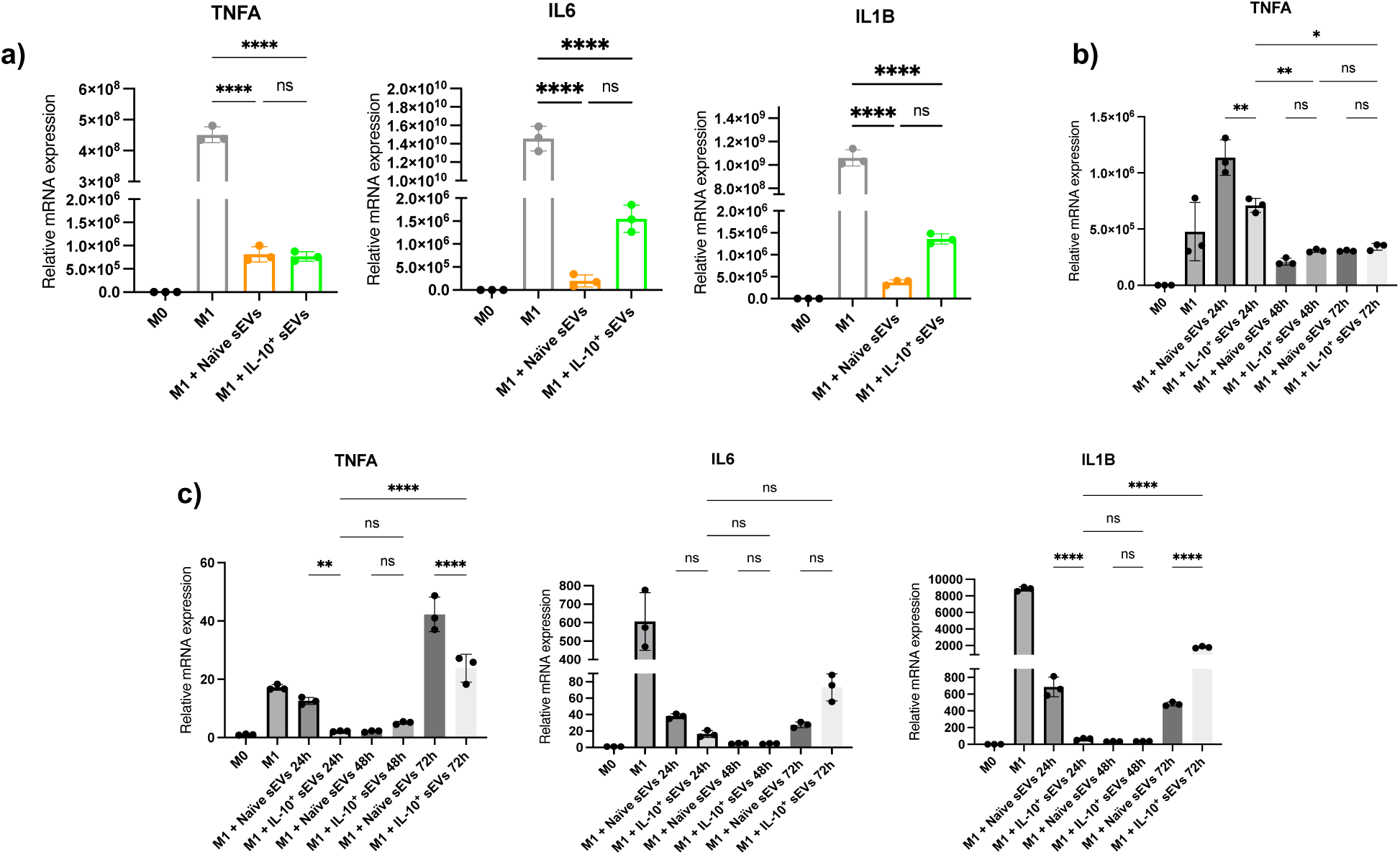
Effect of IL-10^+^ sEVs on inflammatory macrophages. qPCR analysis of mRNA expression levels of pro-inflammatory markers in LPS-stimulated **(a)** human primary macrophages from peripheral blood mononuclear cells (PBMCs), **(b)** RAW 264.7, and **(c)** THP-1 macrophages following incubation with sEVs (10 µg total protein) for 24 h **(a)** or 24 h, 48 h, and 72 h **(b-c)**.

**Supplementary Figure 7.**
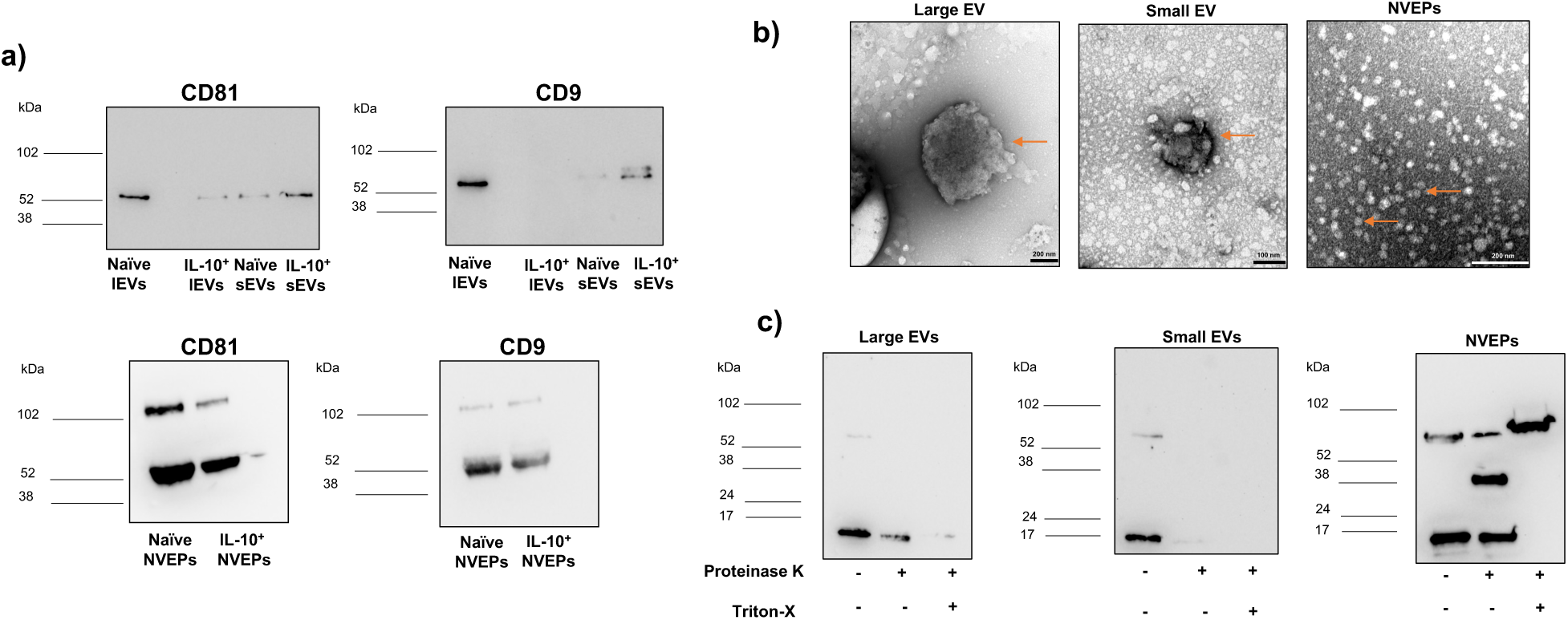
Characterization of EVs and non-vesicular extracellular particles (NVEPs). **(a)** Western blot analysis showing expression of EV markers, CD81 and CD9, in EVs and NVEPs. **(b)** Representative transmission electron microscopy images of EVs and NVEPs (orange arrows) **(c)** Proteinase K protection assay characterizing IL-10 protein in IL-10^+^ EVs/NVEPs, analyzed by Western blotting.

**Supplementary Figure 8.**
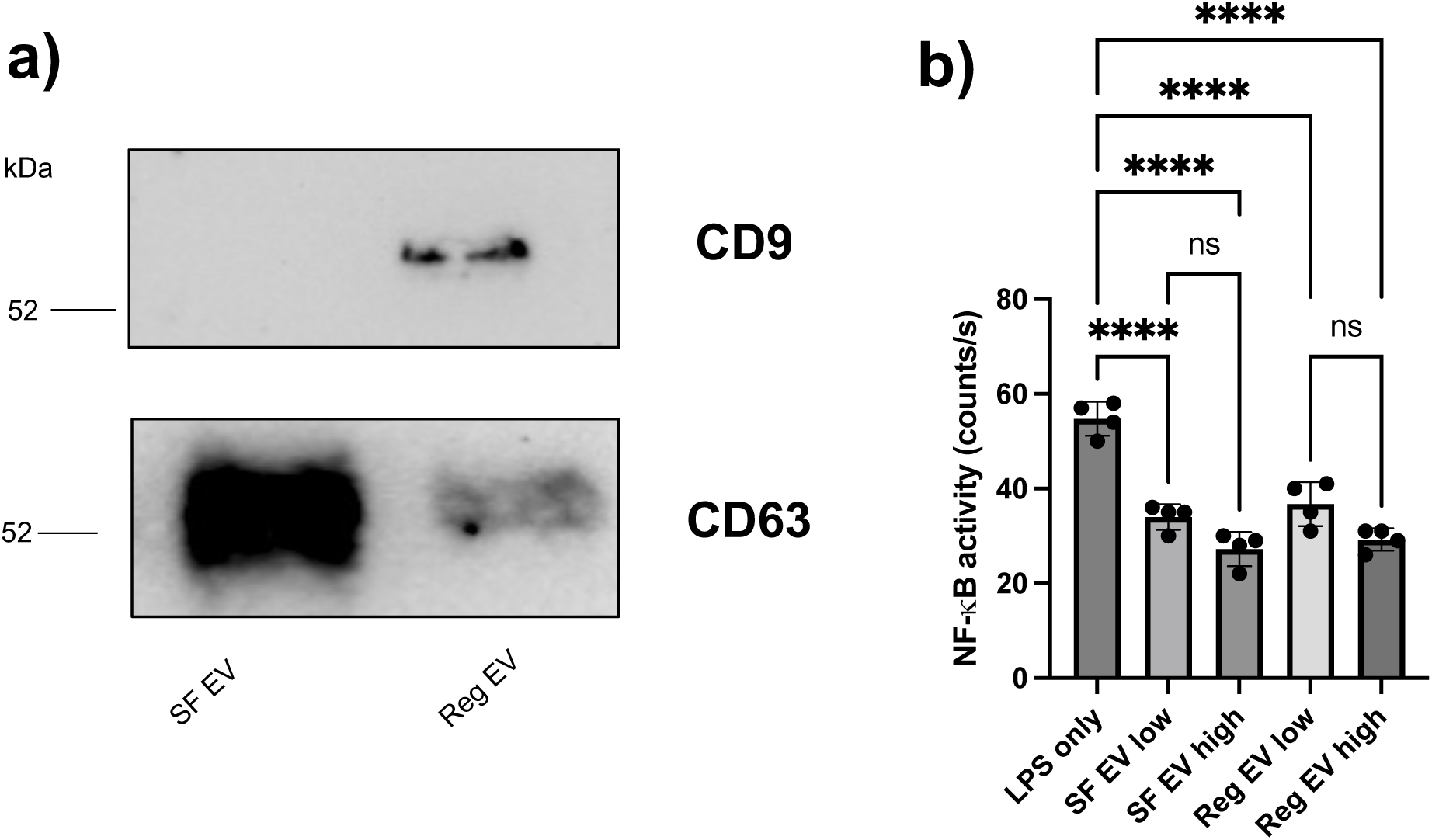
Characterization of sEVs isolated from serum-free versus EV-depleted media condition. **a)** Western blot analysis of CD9 and CD63 in sEVs isolated from serum-free (SF) and EV-depleted (Reg) conditioned media. **b)** NF-κB activity in monocytes measured by bioluminescence assay following incubation with SF or Reg sEVs at low (2 µg) or high (20 µg) protein concentrations.

